# Single-cell proteomics differentiates Arabidopsis root cell types

**DOI:** 10.1101/2024.04.09.588771

**Authors:** Christian Montes, Jingyuan Zhang, Trevor M. Nolan, Justin W. Walley

## Abstract

Recently, major advances have enabled the exploration of cellular heterogeneity using single-cell proteomics. Here we examine the feasibility of single-cell proteomics on plant samples. We focus on *Arabidopsis thaliana*, examining isolated single cells from the cortex and endodermis, which are two adjacent root cell-types derived from a common stem cell. From 756 cells we identify 3,763 proteins and 1,118 proteins/cell. Ultimately, we focus on 3,217 proteins quantified following stringent filtering. Of these, we identified 596 proteins whose expression is enriched in either the cortex or endodermis and are able to differentiate these closely related plant cell-types. Collectivity, our findings underscore the promise of single-cell proteomics to explore the heterogeneity of expression between individual plant cells.

## Introduction

Understanding the coordination of signaling networks across different cell, tissue, and/or organ types is critical for understanding organismal function. Traditional omics profiling methods such as transcriptomics, proteomics, and metabolomics have been instrumental in revealing the differential accumulation of gene products and molecular signaling networks. Yet, these bulk analyses merely capture the mean expression within a mix of cells and cell types, thus concealing the intrinsic heterogeneity of expression between individual cells. Advances in single-cell RNA sequencing (scRNA-seq) are unraveling cellular heterogeneity and providing unique insight into biological systems ^1,2^. It has long been recognized that genome-wide correlations between the levels of proteins and mRNAs are weakly positive in all organisms studied, including plants ^3–6^. Measurement of both mRNA and protein levels in bulk samples gives a more complete picture of cellular state and improves gene regulatory network predictions for bulk tissue samples ^6^. Thus, there has been a concerted effort to develop single-cell proteomics to enable the measurement of not only mRNA but also protein levels from single cells.

Single-cell proteomics has made major gains in both proteome coverage and sample throughput in the last several years ^7–9^. Recent developments have focused on several areas. These include reducing sample preparation volumes to reduce sample loss ^10–17^, ultra-low liquid chromatography (LC) flowrates (<100 nL/min), and optimized LC ^11,12,14,18,15^. Additionally, both label-free and multiplex methods are continually improving proteome coverage and even enabling detection of post-translational modifications ^12,19–23^. Multiplex sample processing has the advantage of enabling greater throughput and a lower amount of instrument time per cell for both Data Dependent Acquisition (DDA) and Data Independent Acquisition (DIA) methods ^24,8,25^.

Recent advancements in single-cell proteomics methodologies have shown considerable promise, yet the application of single-cell proteomics to plants introduces specific challenges. One of the most common methods for obtaining single cells in both animal and plant systems is Fluorescence Activated Cell Sorting (FACS). Plant tissues introduce additional challenges as plant cells are surrounded by cell walls, which needs to be removed from the membrane-encapsulated cell before the resulting protoplasts can be sorted using FACS or other methods ^26,9^. Plants also produce an array of substances such as phenolics, terpenes, various pigments, organic acids, lipids, and polysaccharides which complicate proteomic analysis ^27^. Here, we assess the feasibility of multiplexed single-cell proteomics for plants. We examine single cells isolated from the cortex and endodermis of Arabidopsis roots and show that single-cell proteomics can differentiate the proteomes of these two cell types.

## Experimental Procedures

### Plant material and single cell isolation

*Arabidopsis thaliana* accession Col-0 plants expressing a previously described *pCASP2:NLS:GFP:GUS* reporter for endodermis (*pCASP2:GFP*) ^28^, or a *pCORTEX:erGFP* reporter for cortex (*pCORTEX:GFP)* ^29^, were used for these experiments. Plant growth and protoplast isolation were performed as described ^2^, with minor modifications to accommodate proteomic analysis. Seeds were sterilized using 50% (v/v) bleach with 0.05% Tween-20 for 10-15 minutes. Approximately 2000-3000 seeds per plate were then plated across two rows on 1/2 Linsmaier and Skoog (LSP03-1LT, Caisson Labs; pH 5.7), 1% sucrose media topped with 100 μm nylon mesh (Nitex 03-100/44) to facilitate root collection. The seeds were stratified for 2 days at 4°C in the dark. The plates were then placed vertically in a Percival growth chamber set to 22°C, 16 hours light/8 hours dark, and grown for 5 days. Approximately 0.5-1 cm root tips were harvested with a razor blade and placed into a 35 mm petri dish containing a 70 μm cell strainer and 4.5 mL enzyme solution (1.5% [w/v] cellulase [ONOZUKA R-10, GoldBio], 0.1% Pectolyase [Sigma P3026], 0.4 M mannitol, 20 mM MES (pH 5.7), 20 mM KCl, 10 mM CaCl2). Digestion was allowed to proceed for one hour at 25°C in the dark, shaking at 80 rpm on an orbital shaker. The resulting protoplast solution was filtered twice through 40 µM filters, centrifuged for 500g, 5 minutes, and washed with 2 mL of washing solution (0.4 M mannitol, 20 mM MES (pH 5.7), 20 mM KCl, 10 mM CaCl2). After a second centrifugation of 500g for 3 minutes, protoplasts were resuspended in 1 mL of washing solution and 4′,6-diamidino-2-phenylindole (DAPI) was added to a 14 µM final concentration before fluorescence-activated cell sorting (FACS).

FACS was performed on a Beckman Coulter MoFlo Astrios EQ High-Speed Sorter using a 100 µm nozzle at 28 psi. Live cells for the indicated cell type were sorted using a GFP-positive, DAPI-negative gating strategy. GFP was excited with a 488 nm laser line and acquired with a 526/52 nm filter, and DAPI was excited with a 405 nm laser line and acquired with a 448/59 nm filter. For single-cell collection, one cell per well was sorted into 384 well plates (ThermoFisher AB1384) containing 1 µL of Optima LC-MS/MS grade water (Fischer Scientific W6). Following collection, the plates were sealed with Adhesive PCR plate foil (ThermoFisher AB0626) and immediately frozen on dry ice. For bulk sorting of the carrier and reference samples, the same FACS settings were used, except that cells were sorted into 1.5 mL low-bind tubes containing 50 µL Optima LC-MS/MS grade water.

### Carrier, reference, and 10x Master sample preparation

Carrier and reference samples were processed using a modification of the Filter-Assisted Sample Preparation (FASP) ^30,27^ protocols. In order to make a “10x Master” testing set, tubes containing 52,000 frozen isolated protoplasts from endodermal cells, and 143,000 from cortical cells were incubated at 95℃ for five minutes, followed by water bath sonication for five minutes. Samples were flash frozen in liquid nitrogen and a second round of incubation and sonication was performed. UA buffer (8M urea in 100mM TRIS-HCl pH 8.0) was then added to each tube to a total volume of 750 µL, and thoroughly mixed. Samples were transferred to a Amicon® Ultra Centrifugal Filter, 30 kDa MWCO column (Millipore UFC5030) in three batches of 250 µL each. Columns were centrifuged at 12,000xg until all loading volume was collected (between 10 and 15 minutes). After all sample was loaded onto the column, three washes of 300 µL UA buffer, followed by 2 washes using 100 µL of UA buffer was performed. Finally, the column was conditioned two times with 100 µL of 83.3 mM TEAB, pH 8.5 (Invitrogen). Columns were centrifugated at 12,000xg for 10 minutes between each step. After the last conditioning step, 5.5 µL of enzyme solution (275 ng of Trypsin Gold, Promega V5280; and 8 units of Benzonase, Merck Millipore 70746; in 83.3 mM TEAB pH 8.5) was added to the 27 µL of residual column volume and incubated overnight at 37℃. Samples were then eluted from columns two times using 100 µL of Optima grade H2O (Fischer). Eluted peptides were vacuum-dried in a SpeedVac (Eppendorf), resuspended in 30 µL of Optima grade H2O, and quantified on a Little Lunatic system (Unchained Labs) using the “MS peptide” quantification function ^31^. Peptides from endodermis and cortex cells were combined in equal parts for a total of 550 ng, which corresponds to approximately 3,244 cells, assuming 170 pg of peptides per cell as we previously reported ^9^. Isobaric labeling was done using 18-plex TMTpro reagents (Thermo Scientific A52047) as follows: 2,000 carrier cells (340 ng) resuspended in 2 µL of 83.3 mM TEAB, pH 8.5 were mixed with 1 µL of anhydrous acetonitrile (CAN, Merck AX0143-7) containing 24 ug of 126 TMTpro label; 50 reference cells (8.5 ng) resuspended in 1 µL of 83.3 mM TEAB, pH 8.5 were mixed with 0.5 µL of anhydrous ACN containing 600 ng of 127N TMTpro label; mock single-cell samples (1.7 ng) were made by resuspending 24 ng of cells into 14 µL of 83.3 mM TEAB, pH 8.5 and then split into 14 tubes with 1 µL each. Each tube received 0.5 µL of anhydrous ACN containing 96 ng of one label from 128C to 135N. All samples were then water bath sonicated for 5 minutes, spun down and incubated for 4 hours at room temperature. Label efficiency was confirmed to be ∼99%, then all samples were quenched using 1% hydroxylamine to a final concentration of 0.5%, incubated for 30 minutes, and pooled to create the 10x Master test set.

For carrier and reference samples used in the actual single-cell experiment, 47,000 cells from each cell type were subjected to the same sample preparation protocol. Carrier and reference samples where then labeled with TMTpro 126 and TMTPro 127N, respectively at the same proportion of peptide to label as described above for the 10x Master set. Once sample preparation was done, carrier samples were diluted to 135 cells / µL and reference samples to 5 cells / µL. Samples were stored at -80℃ until single cell samples were ready.

### Single-cell sample preparation

For each cell type, a 384-well plate harboring 1 cell in 1µL of Optima LC-MS/MS grade water per well was used. Samples were processed one plate at a time, as suggested by the SCoPE2 protocol ^20,32^. The previously sealed plate was set on liquid nitrogen for 5 minutes and immediately incubated at 90℃ on an OT-2 temperature module for 10 minutes, and then immediately cooled to 12℃. After incubation, the plate was briefly spun down and water bath sonicated for 5 minutes. This procedure was repeated one more time. The plate sealing film was replaced, the plate was placed on the OT-2 temperature module, and then cooled down to 4℃. Once the plate was cooled, 1 µL of enzyme solution (10 ng of Gold Trypsin and 0.2 units of benzonase in 83.3 mM TEAB, pH 8.5) was dispensed to each well by the robot. Once dispensing was finished, sealing film was replaced and plate was incubated at 37℃ for 3 hours. After incubation, the plate was spun down, placed back on the temperature module, and cooled down to 4℃. Afterwards, Aliquots of TMTpro reagent resuspended in anhydrous ACN at a concentration of 6.47 µg/µL were put in the temperature module at 10℃. To reduce ACN leaking to a minimum, sample plate height was leveled to that of the robot tip when dispensing acetonitrile by setting the plate on the OT-2 magnetic module and both modules were placed right next to each other (i.e., OT-2 deck slots 4 and 7, respectively). Additionally, tip was pre-soaked in ACN by pipetting up/down in the label tube before dispensing and 1µL of TMT label was added to each well. Label assignment was set by randomizing which label each sample would receive while maintaining 7 samples of each cell type per multiplex (Supplemental Table 1). TMTpro labels 127C and 128N were not used. The label map was passed to the OT-2 for dispensing (see “https://github.com/chrisfmontes/Ath_root_SCproteomics” for scripts). Once all wells received label, a new plate seal was added. Samples were water bath sonicated for 5 minutes, vortexed for 5 minutes, spun down, and incubated at room temperature for 2 hours. The labeling reaction was then stopped by addition of 1 µL of 0.25% hydroxylamine to each well. The plate was then sealed with new sealing film, vortexed, spun down, and incubated at room temperature for 30 minutes. Once both plates were fully processed, samples were transferred to 250 µL glass vial inserts (Thermo 6PME03C1SP) and pooled into each multiplex by following the labeling map (Supplemental Table 1). One µL of labeled carrier (135 cells), and 1µL of labeled reference (5 cells) were added to each pooled multiplex. All 54 multiplexes were SpeedVac’d to almost dry and resuspended in 10.2 µL of 0.1% formic acid and 10 µL were injected for each LC-MS run.

### LC-MS/MS

A U3000 HPLC directly coupled using a Nanospray Flex ion source to Q-Exactive Plus high-resolution quadrupole Orbitrap mass spectrometer (Thermo Scientific) was used for LC-MS/MS. For single-cell proteomics, peptides were loaded onto a μPAC Trapping Column at 10 µL min^-1^ and then separated using a 200 cm µPAC analytical column with a gradient optimized using GradientOptimizer ^33^ (Supplemental Table 2) and a flow rate of 300 nL min^-1^. Eluted peptides were analyzed by Data-Dependent Acquisition using Xcalibur 4.0 software in positive ion mode with a spray voltage of 2.20 kV and a capillary temperature of 275 °C and an RF of 60. MS1 spectra were measured at a resolution of 70,000, an automatic gain control of 1e6 with a maximum ion time of 100 ms, and a mass range of 450 to 1600 m/z. Up to 7 MS2 were triggered at a resolution of 70,000 with a fixed first mass of 110 m/z. An automatic gain control of 5e4 with a maximum ion time of 300 ms, an isolation window of 0.7 m/z with 0.3 m/z offset, and a normalized collision energy of 33. Charge exclusion was set to unassigned, 1, 4–8, and >8. MS1 that triggered MS2 scans were dynamically excluded for 30 s.

The raw spectra were analyzed using MaxQuant version 2.4.2.0 ^34^ against the Tair10 proteome file containing 35,386 proteins entitled “TAIR10_pep_20101214” that was downloaded from the TAIR website (https://www.arabidopsis.org/download_files/Proteins/TAIR10_protein_lists/TAIR10_pep_20101214) and was complemented with reverse decoy sequences and common contaminants by MaxQuant. Methionine oxidation and protein N-terminal acetylation were set as variable modifications. The sample type was set to “Reporter Ion MS2” with “18plex TMT selected for both lysine and N-termini” and including impurity corrections for TMTpro lot number XE350091 (126-134N) and XF347836 (134C and 135N). Digestion parameters were set to “specific” and “Trypsin/P;LysC.” Up to two missed cleavages were allowed. No PSM, peptide or protein FDR filtering were used in MaxQuant. Data were further processed using DART-ID ^35^ and the SCP R package, which included protein FDR filtering to < 1% ^36^.

### Data analysis

Statistical analysis to determine differential protein accumulation was assessed by Welch’s two-sample t-test (two-sided) with Benjamini-Hochberg p-value adjustment, on the “rstatix” R package. Principle Component Analysis (PCA) and Uniform Manifold Approximation and Projection (UMAP) were carried out using the “scater” R package. Hierarchical clustering was calculated using the ‘hclust’ base R function to obtain Spearman-based distances, and the package “dendsort” was used to further organize the order of each leaf. Heatmap visualization was performed using the “ComplexHeatmap” R package. See “https://github.com/chrisfmontes/Ath_root_SCproteomics” for the full set of scripts.

## Results and Discussion

The Arabidopsis root represents a well-characterized organ with defined cell types, making it an excellent system for benchmarking single-cell proteomics in plants. We choose to focus on two cell types, the cortex and endodermis, which are derived from a common stem cell ^37,38^. These two cell types are adjacent to each other, offering the opportunity to determine if two proximal plant cell types can be differentiated using SCoPE2 ^20,32^ single-cell proteomics (Figure 1). Plants carrying either *pCASP2:GFP* (endodermis) or *pCORTEX:GFP* (cortex) markers were grown for five days at which point the roots were subjected to enzymolysis to release protoplasts. Endodermis or cortex protoplasts were then separately isolated by FACS. For single-cell collection, a pilot experiment was carried out, where the ability of the MoFlo Astrios EQ High-Speed Sorter to deliver a single cell per well was verified by microscopy. 384 protoplasts from each cell-type were also deposited one per well into 384 well plates. Additionally, for reference and carrier, equal numbers of endodermis and cortex protoplasts were collected.

**Figure 1.**
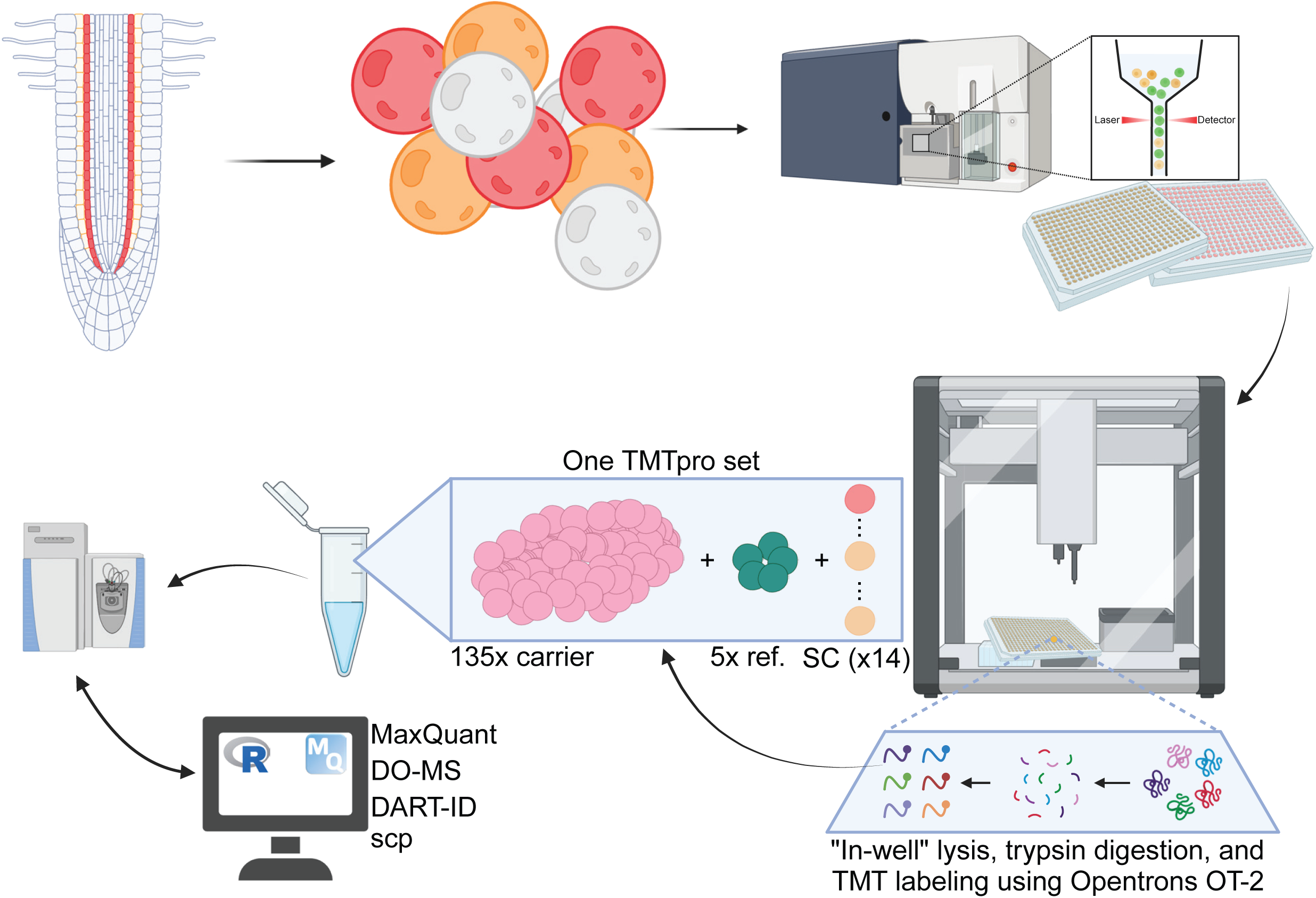
Arabidopsis endodermis and cortex single-cell proteomics pipeline. Single cells (protoplasts) from plants carrying either pCASP2:GFP (endodermis) or pCORTEX:GFP (cortex) markers are isolated by FACS. Samples are processed using on Opentrons OT-2. 54 multiplexed sets consisting of single-cells, reference, and carrier were analyzed by LC-MS/MS.

Single-cell samples were processed based on the Minimal ProteOmic sample Preparation (mPOP) method ^39^ modified to use an Opentrons OT-2 liquid handling robot ^14^. Plants synthesize a wide range of metabolites, which introduces additional challenges during proteomic analysis. We reasoned that the carrier and reference samples would be the major source of interfering compounds. Thus, we processed endodermis and cortex bulk collected protoplasts with a modified urea-FASP method ^27^. We previously determined that Arabidopsis root protoplasts contain on average ∼170 pg of protein ^9^. The ratio of carrier channel to single-cell ratio is an important consideration. Thus, based on previous studies using similar mass analyzers we created 5x reference and 135x carrier samples ^20,40^. Single-cell (128C to 135N), reference (127N), and carrier (126) samples were labeled with TMTpro 18-plex reagents (TMTpro 127C and 128N channels were not used). TMTpro labeled samples were pooled into 54 multiplexed sets, using the OT-2 liquid handling robot, where each set contained 14 single cells, one 5x reference, and one 135x carrier.

Prior to running experimental samples, we created “1x master” test sets containing TMTpro labeled peptides diluted to single-cell equivalent, 5x reference, and 200x carrier samples. Using these test sets, we examined different Q Exactive Plus acquisition settings ^40^. Following MaxQuant spectral searches, we analyzed the resulting data using Data-driven Alignment of Retention Times for Identification (DART-ID) ^35^ and Data-driven Optimization of MS (DO-MS) ^41^. Subsequent evaluation of multiple “1x master” injections led us to conclude that the mass spectrometry parameters described in the original SCoPE2 protocol were optimal for our instrumentation. However, since we used a different analytical column, we optimized the LC gradient using GradientOptimizer ^33^ (Supplemental Table 2). Using the best-performing settings we then analyzed 756 single cells across the 54 multiplexed sets using a 110-minute active (150 minutes total) non-linear LC gradient. The resulting raw data were searched initially using MaxQuant, and further processed using DART-ID ^35^ and the SCP R package ^36^. We assessed the sensitivity and consistency of our data as recommended by Vanderaa and Gatto ^42^ (Figure 2). These assessments suggest our dataset is similar to previous single-cell studies in animal models, including the original SCoPE2 report ^20^. Using initial filters including Precursor Intensity Fraction (PIF), Sample to Carrier Ratio (SCR), and protein & peptide FDR < 1%, we identified 3,763 proteins with an average of 1,118 proteins per cell (Table 1).

**Figure 2.**
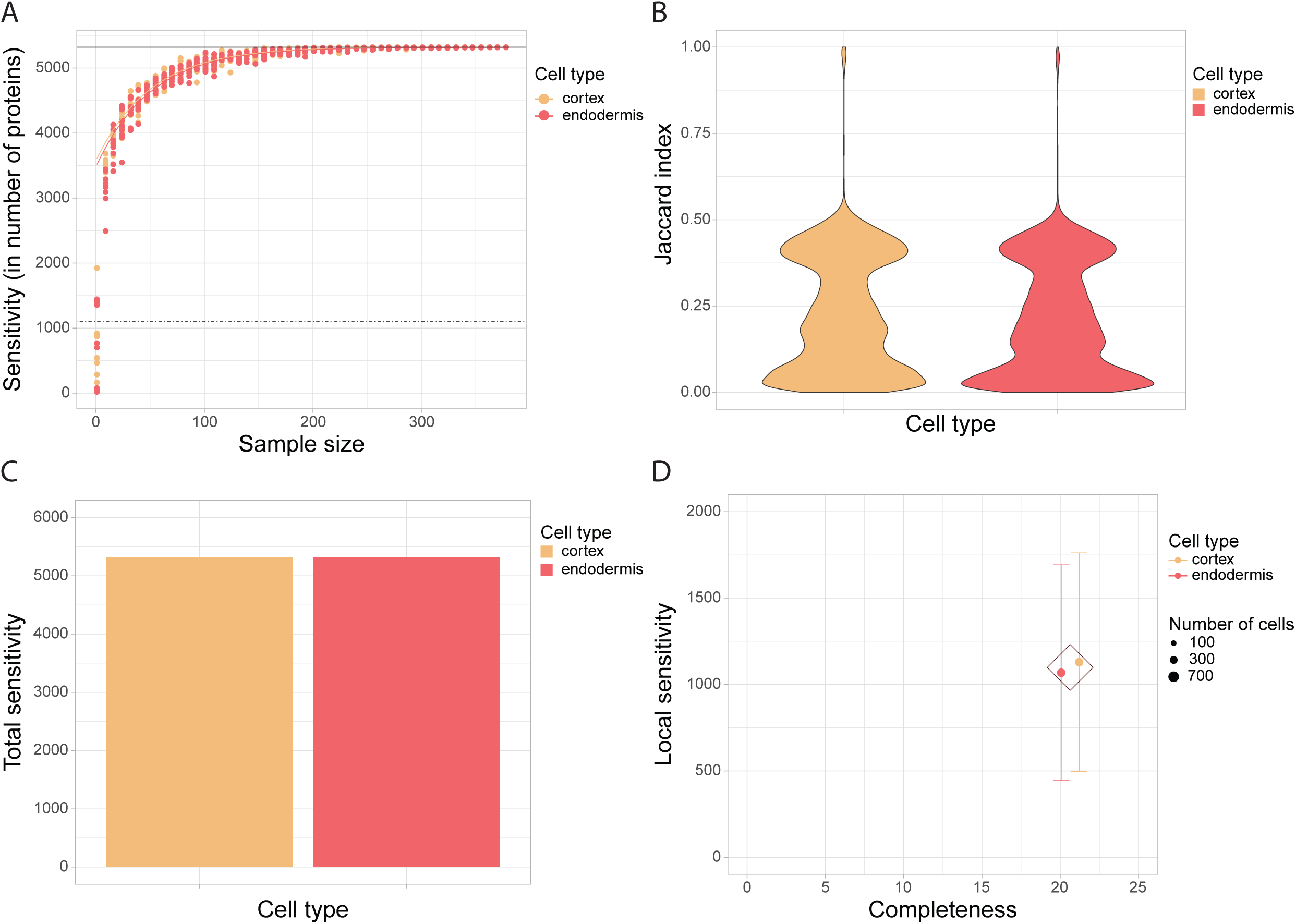
Sensitivity assessment of single-cell proteomics data. A) Cumulative sensitivity curve for both cell types. Dashed line represents average local sensitivity while solid line marks average total sensitivity. B) Violin plot showing Jaccard index per cell type. C) Bar plot showing total sensitivity per cell type. D) Local sensitivity plot. Center dot represents local sensitivity per cell type and bars represent standard deviation. Diamond represents average local sensitivity for both cell types.

**Table 1.** Data summary following each filtering step.

|  | **PSMs** | **NR peptides** | **Proteins** | **Proteins/cell** | **DE proteins** |
| --- | --- | --- | --- | --- | --- |
| **PIF > 0.8** | 186,471 |  |  |  |  |
| **mean SCR < 0.4** | 170,243 |  |  |  |  |
| **Protein FDR < 1%** | 144,965 | 12,323 | 3,763 | 1,118 |  |
| **Cells with median CV < 0.69** | 121,849 | 11,699 | 3,714 | 1,129 |  |
| **Peptides with <99% missing values** |  | 10,393 | 3,542 | 741 |  |
| **Proteins with <98% missing values** |  | 10,393 | 3,217 | 737 | 596 |

We added additional filters, removing cells with extreme peptide Coefficient of Variation (CV) and proteins quantified in less than 98% of the cells to generate a stringent filtered list. While this stringent filtering reduced the number of cells to 81 (33 endodermis cells and 48 cortex), we were able to quantify 3,217 proteins across these cells (Table 1, Supplemental Table 3A). Cells were mainly eliminated due to CV filtering. We speculate that this may be due to differences in root cell size (total protein/cell) as there is a gradient of cell size along the length of the root. Additionally, cells in many plant tissues, including roots, undergo genome endoreduplication ^43,44^. While this generally results in larger cells, the increase in cell size is non-linear, which suggests that identifying normalization factors other than histone proteins ^45^ will be necessary for plant single-cell proteomic studies. This issue of cell size variability potentially contributing to quantitative variation can also be studied in the future using instruments able to sort based not only on fluorescence but also cell size/morphology, such as the BD FACSDiscover S8 Cell Sorter or Cellenion’s cellenONE.

We next explored differences in the proteomes of endodermis and cortex cells for the stringently filtered set. Principal Component Analysis (PCA) and Uniform Manifold Approximation and Projection (UMAP) of the top 25% most variable proteins show a clear separation between endodermis and cortex cells (Figure 3). We also performed differential expression analysis and identified 596 proteins that differentially accumulate in endodermis vs cortex cells (Figure 4A, Supplemental Table 3B-C). We observed enrichment of known proteins in the expected cell-type (Figure 4A). For example, CASP1 an endodermal marker involved of casparian strip (CS) formation ^46,47^, is enriched in the endodermal cells. Additionally, NPY2/MEL3 is enriched in endodermal cells, where it has been reported to control PIN1 subcellular basal localization and root gravitropic response ^48^. We also identified RBOHD enriched in endodermal cells where is known to activate ROS-dependent signaling to help CS formation ^49,50^. Conversely, ARF5 is enriched in the cortex, expression of this gene in the cortex can drive cortical cell division ^51^. At the same time, we found the cochaperone P23-1 enriched in cortex, where it has been reported to be crucial for auxin distribution and proper number of cortical cells ^52,53^. Finally, we found the proteasome 19S regulatory particle 8a (RPN8a) enriched in cortical cells, here it interacts and induces the degradation of BRAHMA to promote boron tolerance ^54^. We also preformed hierarchical clustering on both cell-types (columns) and proteins (rows). Our analysis revealed that cells from the endodermis tend to group together, as do cells from the cortex, indicating a distinct clustering based on cellular type (Figure 4B).

**Figure 3.**
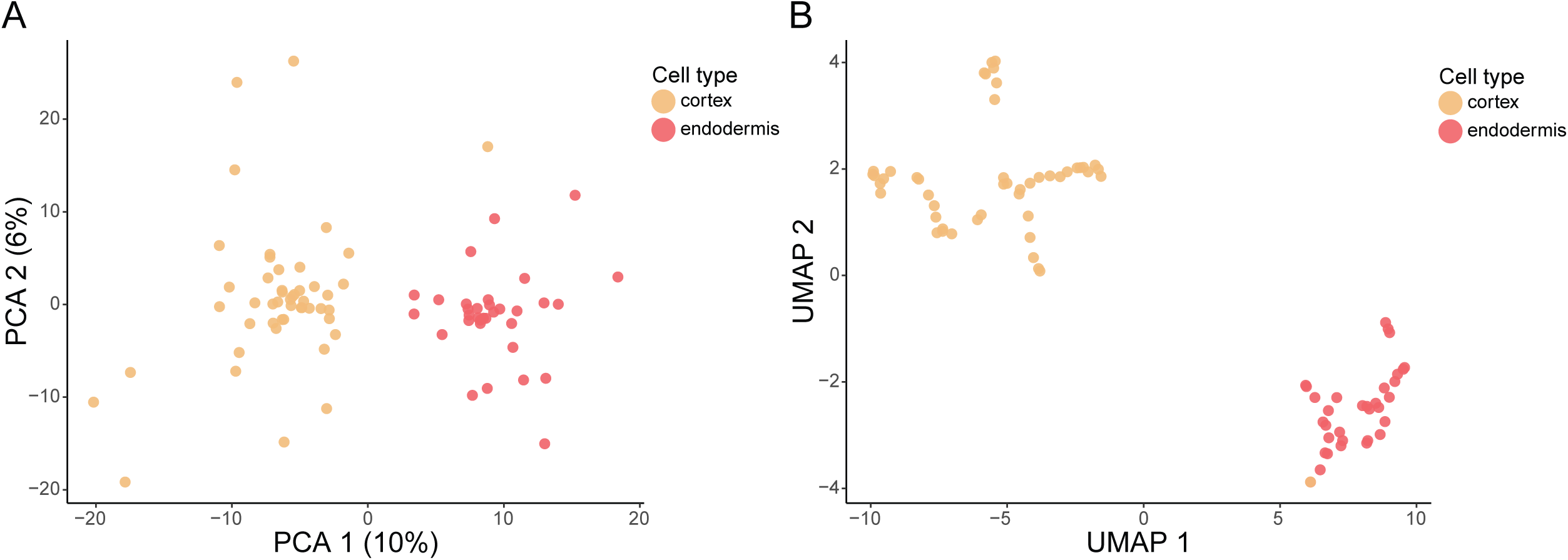
Principal Component Analysis (PCA) and Uniform Manifold Approximation and Projection (UMAP) plots. To calculate PCA and UMAP, the 25% most variable proteins were used.

**Figure 4.**
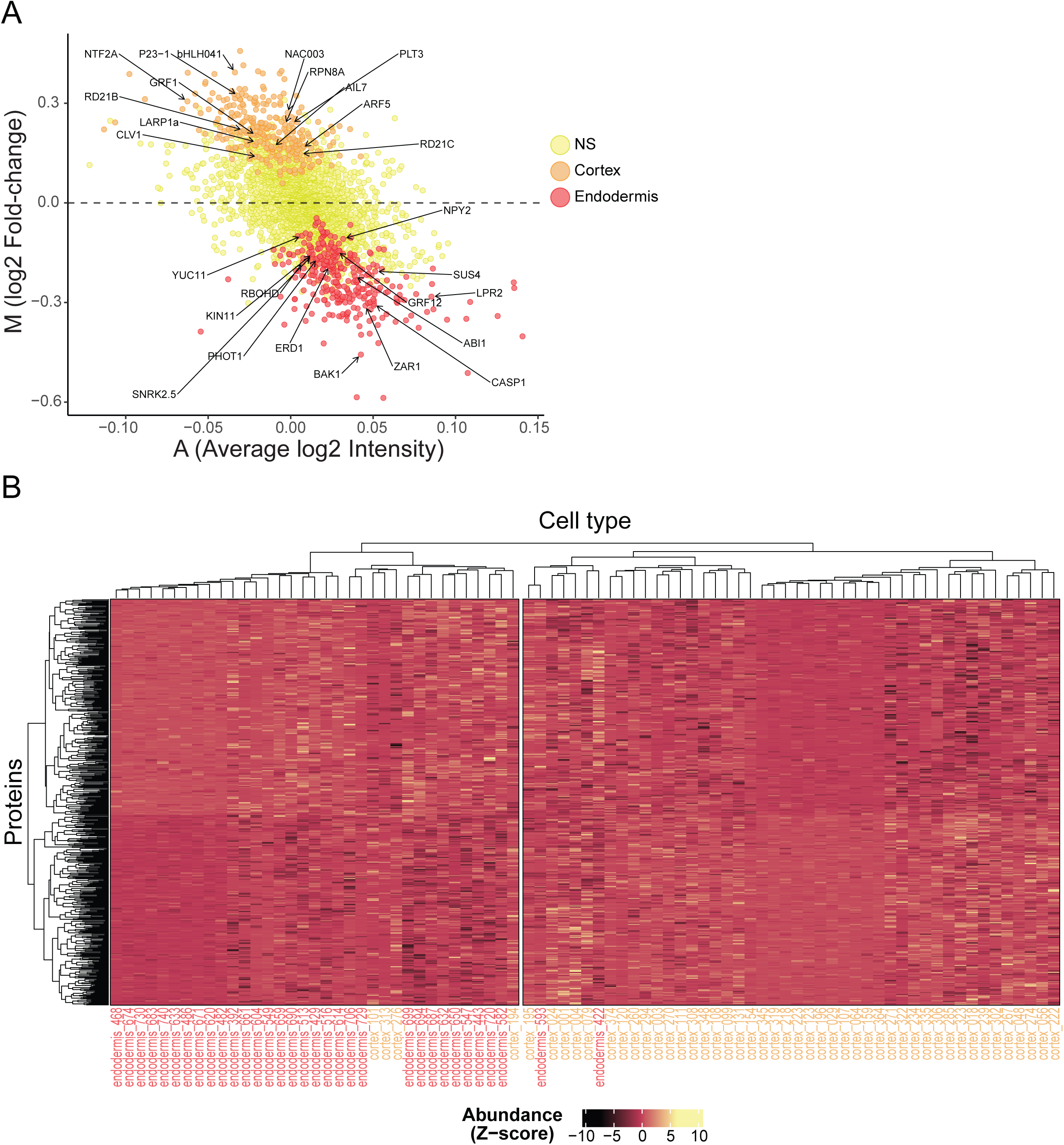
Differential expression analysis. A) MA plot showing proteins enriched in cortical cells (tan), and endodermal cells (pale red). B) Hierarchical cluster and heatmap for the 596 proteins significantly enriched in one cell type (vertical axis) and cells (horizontal axis).

## Conclusions

In recent years, advancements in sample preparation, liquid chromatography, and mass spectrometry have significantly enhanced the capacity of single-cell proteomics, elevating protein detection capabilities from several hundred to several thousand. In the present study we explore single-cell proteomics for examining plant samples. Our results demonstrate the feasibility of plant single-cell proteomics and the ability to differentiate closely related plant cell-types. As with non-plant tissues, future experiments incorporating approaches such as multiplexed DIA ^25,55^ or prioritized SCoPE ^56^ will enable high-throughput quantification of plant single-cell proteomes while limiting missing values inherent in standard DDA approaches. Furthermore, using cell sorting methods that categorize cells by size will clarify how differences in cell size affect measurement variability. This will also assist in pinpointing proteins that could serve as factors for cell size normalization in plant studies. ^45^. Altogether, these results highlight the potential for plant single-cell proteomics to provide unprecedented insights into the cellular heterogeneity of plants and enable a deeper understanding of understanding of the spatiotemporal response to various biological stimuli.

## Acknowledgments

Flow Cytometry was performed by Lynn Martinek in the Duke Cancer Institute Flow Cytometry Facility at Duke University, Durham, NC, which is supported by the NCI Cancer Center Support Grant (CCSG) award number P30CA014236. We dedicate this manuscript to Philip Benfey, who pioneered omics maps of roots.

## Author Contributions

JWW, CM, and TMN planned and designed the research. CM, TMN, JZ performed experiments and/or analyzed data CM, TMN, and JWW wrote the manuscript.

## Data Availability

Proteomics data can be downloaded from MassIVE (http://massive.ucsd.edu) using the identifier MSV000094558.

